# The Incisive Duct as a Pathway for Early Vomeronasal Communication in Neonatal Dogs

**DOI:** 10.1101/2024.06.23.600254

**Authors:** Eva Sanmartín Vázquez, Irene Ortiz-Leal, Mateo V. Torres, Patrycja Kalak, Dominika Kubiak-Nowak, Michał Dzięcioł, Pablo Sanchez-Quinteiro

## Abstract

The detection of chemical signals by the vomeronasal organ (VNO) is critical for communication among mammals from an early age, influencing behaviors such as suckling and recognition of the mother and conspecifics. Located in a concealed position at the base of the nasal cavity, the VNO features a duct covered with a sensory epithelium rich in neuroreceptors. A critical aspect of VNO functionality is the efficient access of stimuli from the nasal and oral cavities to the receptors. In adult dogs, it has been demonstrated through in vivo magnetic resonance imaging and anatomically postmortem how the VNO duct (VD) communicates to the environment through the incisive duct (ID). However, in newborn puppies, the existence of functional communication between the ID and the VD has not been confirmed to date, raising doubts about the potential physiological obliteration of the ID due to its small size and the degree of immaturity of the puppies at birth. Determining this aspect is necessary to evaluate the role played by chemical communication in this critical phase for the survival and socialization of puppies. This study employs serial histological staining techniques to examine the presence and functionality of the incisive duct in neonatal dogs. The serial histological sections have confirmed both the existence of functional communication between both the vomeronasal and incisive ducts in perinatal puppies, and the dual functional communication of the incisive duct with the oral and nasal cavities. The ID shows an uninterrupted lumen along its path and is associated with a sophisticated cartilaginous complex that prevents its collapse, as well as erectile tissue rich in blood vessels and connective tissue that acts as a cushion, facilitating its action under pressure induced by sampling behaviors such as tonguing. This investigation demonstrates the communicative capabilities of the VNO during the perinatal stage in dogs.

## INTRODUCTION

The detection of chemical signals emanating from the environment is an essential part of communication among animals that influences many aspects of their social and sexual behavior, from predator avoidance to reproduction (Pérez-Gómez et al. 2015; Navarro-Moreno et al. 2020). These signals can be detected in the main olfactory system, through receptors located in the back of the nasal cavity (Barrios et al. 2014) or in the vomeronasal system (VNS), which is specialized in the detection of pheromones and kairomones (Bakker and Leinders-Zufall 2016), although there is a significant functional overlap between the two systems (Fortes-Marco et al. 2013; Wyat 2013; Salazar et al. 2016). The VNS is made up of the vomeronasal organ (VNO), where the receptor neurons are located (Torres et al. 2023); the accessory olfactory bulb (AOB), which receives information from the VNO via the vomeronasal nerve (McCoter 1912) and acts as the first integrative center of the VNS (Ortiz-Leal et al. 2022), and the vomeronasal amygdala (Gutiérrez-Castellanos et al. 2014). The VNO is a paired tubular structure located at the base of the nasal cavity, inside of which is the vomeronasal duct (VD), lined on the medial wall by the sensory epithelium containing the neuroreceptors and on the lateral wall by a non-sensory epithelium (Torres et al. 2022). Surrounding the duct are distributed numerous blood vessels, nerves and glands, collectively referred to as the parenchyma, all enclosed by the vomeronasal cartilage (Salazar et al. 1997; Salazar et al. 1998).

Given the location of the sensory neurons within the duct, a fundamental condition for chemical signal detection is the existence of a system that transports the molecules through to the receptors, as well as way of entry from the outside environment. The movement of molecules within the vomeronasal duct is favored by the contraction and dilatation of the parenchymal blood vessels in a mechanism under autonomous control known as the vomeronasal pump (Meredith and O’Connell 1979; Lippner et al. 2024). Furthermore, the existence of an independent control mechanism mediated by parenchymal smooth muscle has been recently proved in mice (Hamacher et al. 2024).

Due to the great topographical and morphological diversity of the vomeronasal organ, communication with the external environment is variable, and two models can be differentiated. In most adult mammals, including domestic ones such as pigs, cows, cats and dogs (Salazar et al. 1995), and wild ones as New World monkeys (Smith et al. 2011), opossums (Poran 1998), hedgehogs (Kondoh et al. 2021), mink (Salazar et al. 1994) and bears (Tomiyasu et al. 2017), it takes place indirectly through the incisive or nasopalatine duct (ID). However, in a smaller number of species, including rodents and lagomorphs, it opens directly into the nasal cavity (Villamayor et al. 2018; Torres et al. 2020; Tomiyasu et al. 2022; Ruiz-Rubio et al. 2023). The incisive duct opens ventrally in the incisive papilla, from where it runs dorsocaudally through the palatine fissure to the floor of the nasal cavity, opening in the ventral meatus (Salazar et al. 2013). Therefore, in adult dogs and wild canids, it provides a means of communication between the nasal and oral cavities as well as access to the vomeronasal duct (Adams and Wiekamp 1984; Dzięcioł et al. 2020; Ortiz-Leal et al. 2024).

Regarding its development in the perinatal stage, although prenatal development of the incisive duct has been proven in a small number of other mammals such as rodents (Coppola and Millar 1994), sheep (Salazar et al. 2003a) and pigs (Salazar et al. 2003b), to our knowledge, the existence of a functional communication in newborn dogs has not been verified. It is necessary to keep in mind that the extrapolation of knowledge concerning the vomeronasal system between different species is problematic due to the great variability that characterizes it (Salazar and Sanchez Quinteiro 2009).

It is surprising that there is no specific information on this aspect, as chemical communication plays a vital role in the early life stages of newborn dogs, serving as a fundamental mechanism for critical behaviors such as recognizing the mother, initiating lactation, and even recognizing their littermates (Kokocińska-Kusiak et al. 2021). The ability to process and respond to these chemical signals is crucial for the development of social behaviors, stress responses, and overall health in the critical neonatal phase (Hepper and Wells 2006).

In this study, we conducted a comprehensive examination of the incisive region in newborn dogs. After completing full histological series in both horizontal and transverse planes, we applied various histological staining techniques to determine whether there is a connection between the vomeronasal organ and the incisive duct, and to assess the functional maturity of the incisive duct. Our findings aim to enhance the understanding of the vomeronasal system in dogs.

## METHODS

For this study, we used a sample of two perinatal dogs obtained from cooperating veterinary clinics. Both puppies died naturally just before birth. After separating the heads, they were preserved immediately in formalin. The nasal cavities of both dogs were decalcified for 24h with Osteomoll (Sigma) while continuously stirring at room temperature. After this process, the samples were embedded in paraffin and cut by a microtome with a section thickness of 7 μm, transversally in one case and horizontally in the other. The resulting histological series were stained using hematoxylin-eosin, alcian blue (AB) and Periodic Acid-Schiff (PAS) stains according to the protocol described in Torres et al. (2021).

Digital images were captured using a Olympus SC180 camera coupled with a Olympus BX50 microscope. An image stitching software, PTGuiPro, was used to obtain images composed of several photographs due to the size of the structures studied. Adobe Photoshop CC was used to adjust brightness, contrast and white balance levels; however, no enhancements, additions or alterations of the image features were made.

## RESULTS

The microscopic study of extensive histological series of transverse sections from the most anterior portion of the nasal cavity and incisive area allowed us to demonstrate the existence of a functional communication between the lumen of the vomeronasal duct and the incisive duct (Fig. 1).

**Figure 1.**
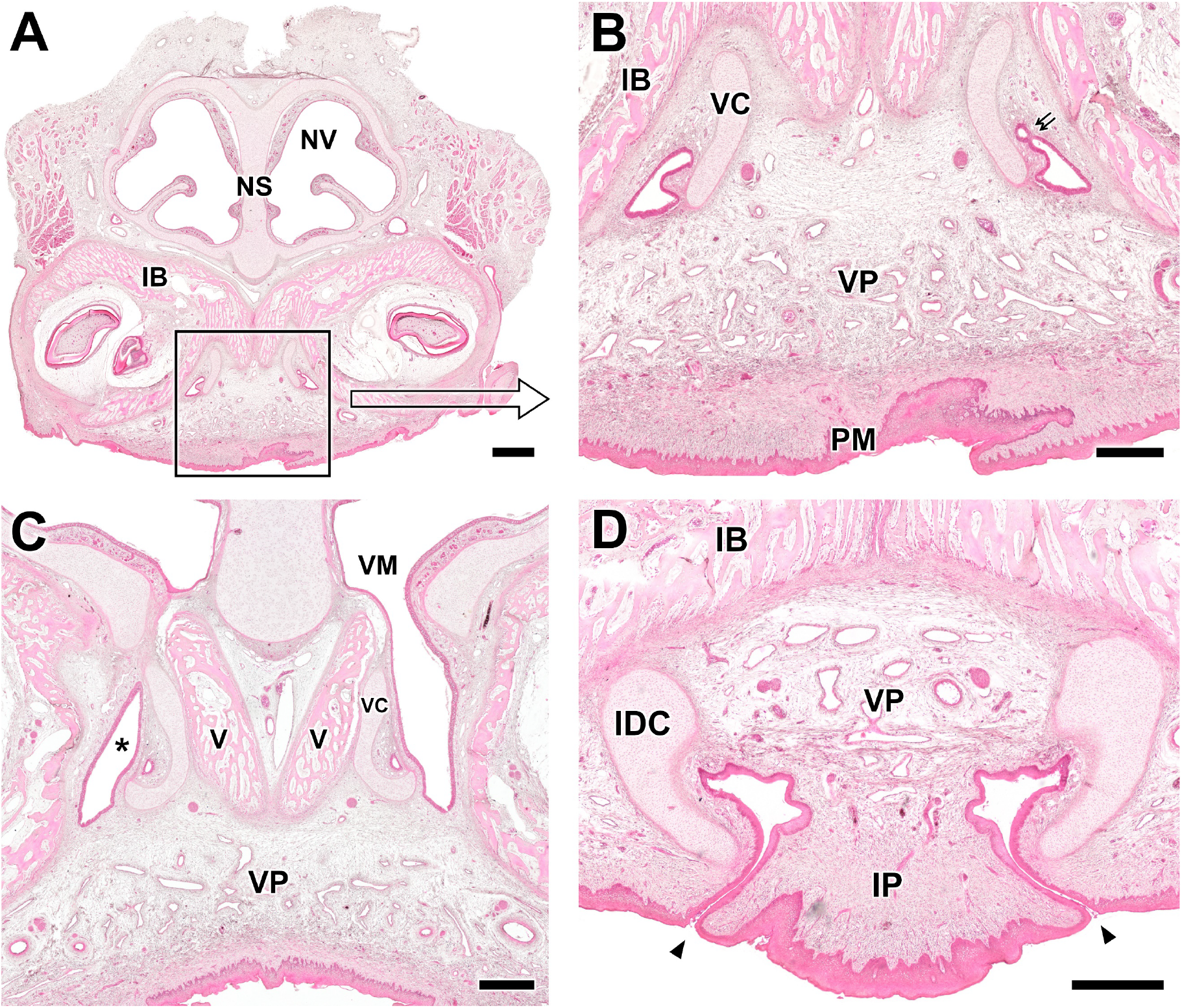
Histological study of the incisive area and rostral portion of the vomeronasal organ of the perinatal dog. **A**. Transverse section of the nasal vestibule (NV) and incisive area at the level where the vomeronasal duct and incisor duct meet. **B**. Close up of the area marked in A, showing the junction point between the vomeronasal and incisive duct (double arrow). The vomeronasal cartilage (VC) is present at this level as a rudimentary plate, which accompanies the medial part of the VNO and incisor duct, while the lateral part is in direct relation to the incisor bone (IB). In the area of the soft palate there is a wide vascular plexus (VP). **C**. Cross caudal section to the previous one showing the vomeronasal organ fully formed. The vomeronasal cartilage rests directly on the two laminae that constitute the vomer bone (V). The incisor canal (*) opens into the nasal cavity at its ventral meatus (VM). **D**. Cross section of the rostral region at the level of the incisive papilla (IP). which shows the existence of a functional communication between the oral cavity and the incisor duct (arrowhead). The incisor duct has a prominent comma-shaped lateral cartilaginous lining (IDC). In the connective tissue between the two incisive ducts a large vascular plexus develops. IT: incisive tooth, PM: palatal mucosa, NS: nasal septum. Hematoxylin-Eosin staining. Scale bar: A = 1 mm, B-D = 500 μm.

This connection is established at a central level of the ID, where both the VD and ID are protected laterally by the incisive bone and medially by the most rostral portion of the vomeronasal cartilage, which arises as a prolongation of the medial part of the incisive duct cartilage (IDC). In more caudal sections, the opening of the ID was observed, in the ventral meatus of the nasal cavity (Fig. 1C). At this point, the VNO is already visible and fully constituted, covered by the J-shaped vomeronasal cartilage. Rostrally, the incisive duct is communicated with the oral cavity through its opening in the incisive papilla (Fig. 1D). At this level, the IDC covers the ducts laterally and a highly developed vascular plexus is located medially.

With the aim of completing the characterization of this area, a complete histological series was also done in the horizontal plane, which confirmed the direct communication between the VD and the ID (Fig. 2). This plane also shows the rich vascular plexus in the medial area between the ducts, which is organized into two different regions: a caudal component with an abundance of blood vessels and a rostral component with a more prominent amount of loose connective tissue.

**Figure 2.**
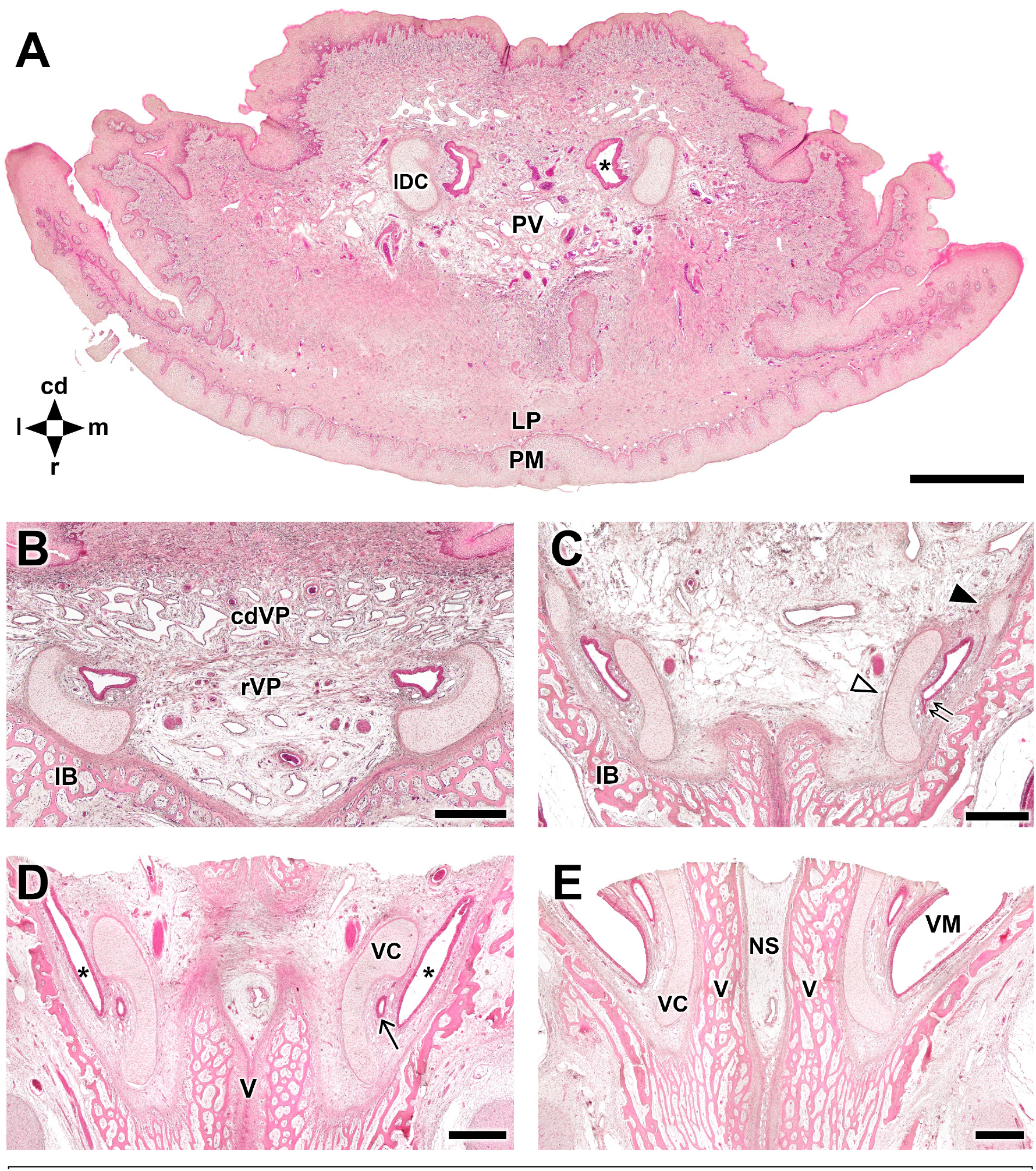
Serial histological study in the horizontal plane of the incisive region. **A**. Ventral section at the level of the incisive papilla showing the presence of broad incisive ducts (*) laterally associated with the cartilage of the incisive duct (IDC). The medial and caudal area to the incisive ducts presents a highly developed vascular plexus (VP). **B**. Section dorsal to the previous one in which two areas of the vascular plexus are differentiated, a rostral one with a greater density of connective tissue (rVP) and another caudal one predominantly with venous plexuses (cdVP). Note the positioning of the cartilage of the incisive duct in direct contact with the incisive bone (IB). **C**. Section dorsal to the previous one, pertaining to the level where the communication between the vomeronasal and incisor ducts is established (double arrow). The incisive duct cartilage is significantly reduced (closed arrowhead) and the ducts are now surrounded medially by the vomeronasal cartilage (open arrowhead). **D**. Section dorsal to the previous one showing the vomeronasal duct (arrow), now independent of the incisor duct (*). The vomeronasal cartilage (VC) covers it medially, being supported by the rostral projection of the vomer bone (V). **E**. Section dorsal to the previous one showing the opening of the incisive duct to the ventral meatus of the nasal cavity (VM). The vomeronasal cartilage keeps direct contact with the vomer bone, which in turn supports the nasal septum (NS). cd: caudal, l: lateral, LP: lamina propria, m: medial, PM: palatal mucosa, r: rostral. Hematoxylin-eosin staining. Scale bars: A= 1 mm, B-E = 500 μm.

Along the dorsoventral axis, the IDC displays an asymmetrical growth pattern of its lateral and medial ends. At the point of the incisive papilla, the cartilage is located laterally to the ID (Fig. 2A) and a slightly more dorsal section shows its direct topographic relation with the incisive bone. However, as it advances dorsally, the lateral end is reduced and the cartilage grows medially, to the point where at the convergence with the VD the ID is nestled between the IDC and incisive bone. This medial sheet of the IDC contributes to the formation of the rostral end of the vomeronasal cartilage.

Finally, in order to characterize the secretions present in the ID, we employed PAS staining, which binds mucopolysaccharides in general, and AB staining, which binds acidic mucopolysaccharides only. Small amounts of PAS-positive and AB-positive secretion were found in the apical part of the epithelium lining the incisive ducts, both in the horizontal and transverse planes (Figs. 3 and 4).

**Figure 3.**
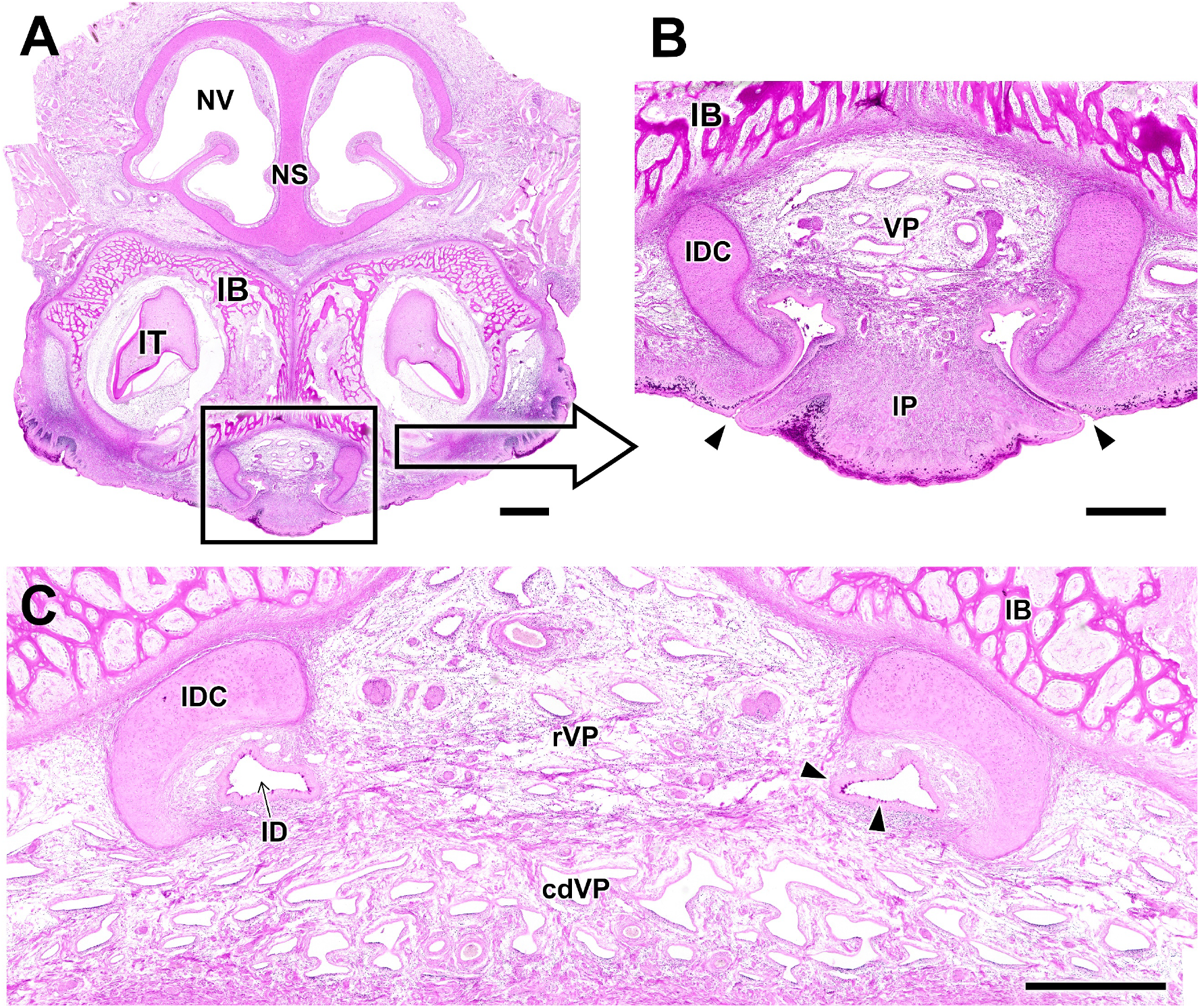
PAS staining of the incisive duct region. **A**. Transverse section of the nasal vestibule (NV) and incisive area, showing a strong staining in the cartilaginous tissue of the nasal septum (NS) as well as the incisive bones (IB) and teeth (IT). **B**. Higher magnification of the area indicated in A, showing the opening of the incisive ducts into the oral cavity (arrowheads) through the incisive papilla (IP). PAS staining reveals the organization of the connective tissue medial to the ducts, featuring a large vascular plexus (VP). **C**. Horizontal section at the level of the incisive papilla, showing the division of the connective tissue in a rostral vascular plexus (rVP) and a caudal vascular plexus (cdVP). The apical part of the epithelium contains a PAS-positive secretion (arrowheads). IDC: incisive duct cartilage. Scale bars: A = 1mm; B, C = 500 μm.

**Figure 4.**
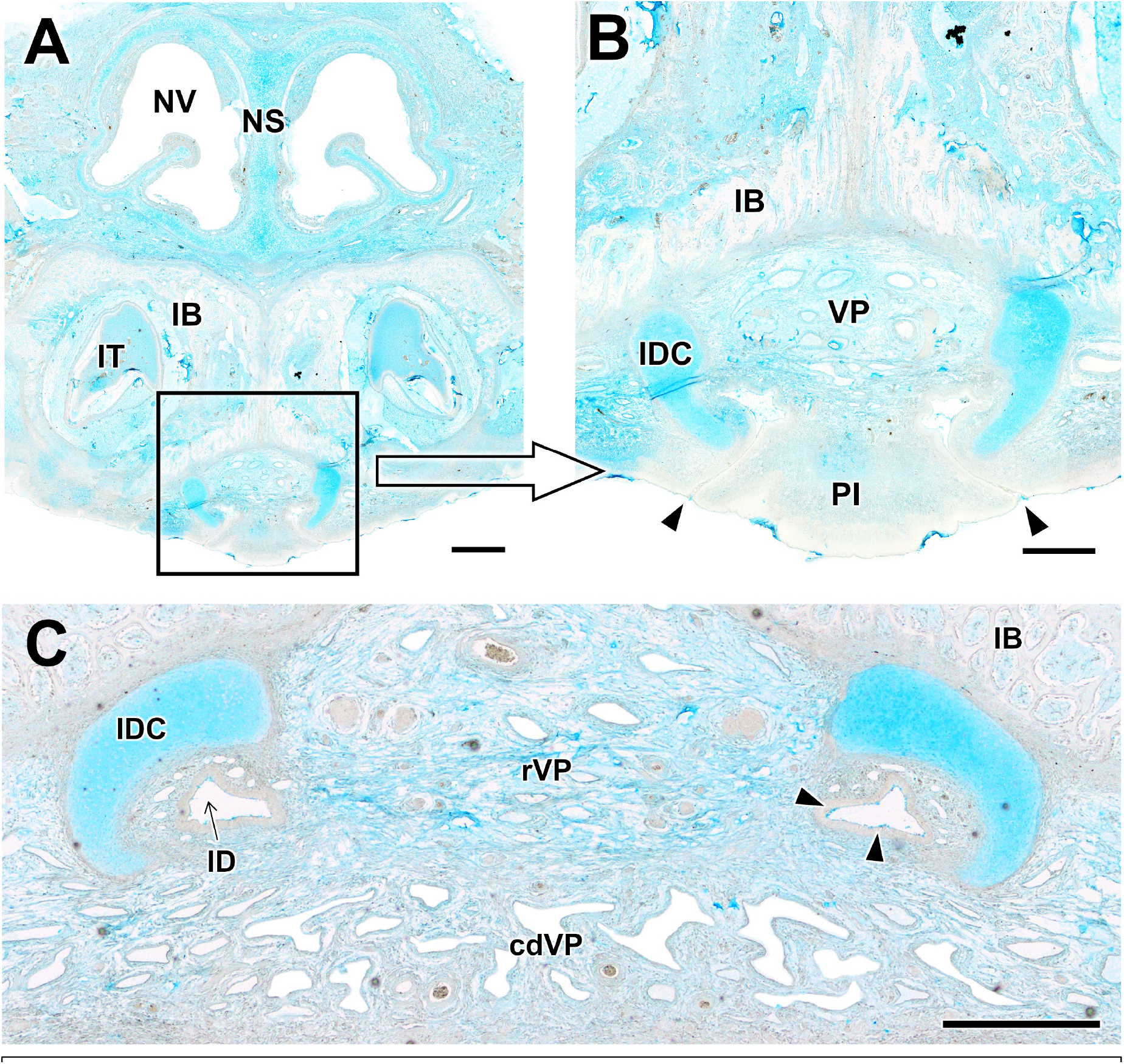
AB staining of the incisive duct region. **A**. Transverse section of the nasal vestibule (NV) and area surrounding the incisive papilla (IP). The mucopolysaccharides in the nasal septum (NS), incisive teeth (IT) and incisive duct cartilages (IDC) are strongly stained by AB. **B**. Higher magnification of the area indicated in A, showing the opening of the incisive ducts into the oral cavity and the vascular plexus medial to the ducts (VP). **C**. Horizontal section showing an AB-positive secretion (arrowheads) in the incisive duct (ID) epithelium. cdVP: caudal vascular plexus, IB: incisive bone, rVP: rostral vascular plexus. Scale bars: A = 1 mm; B, C = 500 μm

Moreover, due to the widespread distribution of mucopolysaccharides in connective tissue, these staining techniques are also useful for the study of its configuration and organization. In the horizontal plane, the distinction between the two areas of connective tissue located rostrally and caudally becomes evident (Figs 3C and 4C).

## DISCUSSION

To our knowledge, the existence of a way of communication between the outside environment and the VNO in dogs in the perinatal stage is an unexplored question. Recently, magnetic resonance imaging provided evidence of in vivo communication in adults (Dzięcioł et al. 2020), but the small size of the structures in the pup hinders the application of this methodology. The execution of extensive histological series in the entire incisive area, both horizontally and transversally, allowed us to solve two fundamental issues: the existence of an effective communication pathway in the newborn dog despite its developmental immaturity, and the model by which it is established.

In our study, we found that there is indeed a functional connection between the vomeronasal duct and the incisive duct, which opens both into the nasal and oral cavities. Moreover, the lumen of the incisive canal is sufficiently wide and clear along its entire length to ensure the entry of the signaling molecules. In addition, the glandular activity observed in the duct epithelium indicates the presence of mucous secretions whose fundamental role in the transfer of semiochemicals to the VNO and in the solubilization of volatile compounds has previously been established (Guiraudie 2003; Lee et al. 2008)

In the region surrounding the incisive canals, we found a vastly developed vascular plexus that provide the tissue with erectile properties. This, along with the presence of the incisive duct cartilages and the incisive bone as a structural reinforcement to the ducts, indicates the presence of a mechanism similar to that of the vomeronasal pump ((Meredith and O’Connell 1979; Hamacher et al. 2024), which promotes the entry and expulsion of the contents of the duct by alternating the contraction and dilatation of the blood vessels, creating a cycle of negative and positive pressure inside the duct (Lippner et al. 2024).

The presence of this vascular, connective, and cartilaginous complex, which is directly linked to the incisive duct, is likely to have a direct connection with the tonguing behaviour mechanism (Liinamo et al. 1997), a specific behavioral mechanism in dogs comparable to the *flehmen* response in cats (Hart 1987). Through repeated pressure on the area of the incisive papilla, its spongy nature would facilitate the movement of fluids towards the interior of the incisive papillation until reaching the outlet point of the nasal duct. At this point, the action of the vasculature would be sufficient to allow for the continuous sampling of the content of the duct.

The current findings on the existence of a functional connection between the vomeronasal organ and the external environment in the perinatal stage confirm the presence of a direct communication between the vomeronasal duct and the external environment in puppies at birth. This suggests the relevant role played by chemical communication in the newborn dog. Additionally, this work provides detailed information on the morphofunctional configuration of the incisive papilla area, which we consider a valuable addition to the study of the vomeronasal system in canids.

## AUTHORS CONTRIBUTION

Conceptualization, E.S.V., I.O.L., M.D., and P.S.Q; Methodology, E.S.V., M.V.T., P.K., D.K., M.D., and P.S.Q; Investigation, E.S.V., I.O.L., M.V.T., P.K., D.K., M.D., and P.S.Q.; Resources, E.S.V., I.O.L., M.D., and P.S.Q; Writing – Original Draft Preparation, E.S.V., I.O.L., and P.S.Q; Writing – Review & Editing, E.S.V., I.O.L., M.D., and P.S.Q; Supervision, E.S.V., I.O.L., M.D., P.S.Q.; Project Administration, P.S.Q.; Funding Acquisition, M.D., P.S.Q.

## COMPLIANCE OF ETHICAL STANDARDS

### Ethical approval

All the animals employed in this study dead by natural causes.

### Informed consent

No human subject was used in this study.

